# Differential gene expression profiling reveals potential biomarkers and pharmacological compounds against SARS-CoV-2: insights from machine learning and bioinformatics approaches

**DOI:** 10.1101/2022.03.30.486356

**Authors:** M. Nazmul Hoque, Md. Arif Khan, Md. Arju Hossain, Md Imran Hasan, Md Habibur Rahman, Mahmoud E. Soliman, Yusha Araf, Chunfu Zheng, Tofazzal Islam

## Abstract

SARS-CoV-2 continues to spread and evolve worldwide, despite intense efforts to develop multiple vaccines and therapeutic options against COVID-19. Moreover, the precise role of SARS-CoV-2 in the pathophysiology of the nasopharyngeal tract (NT) is still unfathomable. Therefore, we used the machine learning methods to analyze 22 RNA-seq datasets from COVID-19 patients (n=8), recovered individuals (n=7), and healthy individuals (n=7) to find disease-related differentially expressed genes (DEGs). In comparison to healthy controls, we found 1960 and 153 DEG signatures in COVID-19 patients and recovered individuals, respectively. We compared dysregulated DEGs to detect critical pathways and gene ontology (GO) connected to COVID-19 comorbidities. In COVID-19 patients, the DEG– miRNA and DEG–transcription factors (TFs) interactions network analysis revealed that E2F1, MAX, EGR1, YY1, and SRF were the most highly expressed TFs, whereas hsa-miR-19b, hsa-miR-495, hsa-miR-340, hsa-miR-101, and hsa-miR-19a were the overexpressed miRNAs. Three chemical agents (Valproic Acid, Alfatoxin B1, and Cyclosporine) were abundant in COVID-19 patients and recovered individuals. Mental retardation, mental deficit, intellectual disability, muscle hypotonia, micrognathism, and cleft palate were the significant diseases associated with COVID-19 by sharing DEGs. Finally, we detected DEGs impacted by SARS-CoV-2 infection and mediated by TFs and miRNA expression, indicating that SARS-CoV-2 infection may contribute to various comorbidities. These pathogenetic findings can provide some crucial insights into the complex interplay between COVID-19 and the recovery stage and support its importance in the therapeutic development strategy to combat against COVID-19 pandemic.

**IMPORTANCE:** Despite it has now been over two years since the beginning of the COVID-19 pandemic, many crucial questions about SARS-CoV-2 infection and the different COVID-19 symptoms it causes remain unresolved. An intriguing question about COVID-19 is how SARS-CoV-2 interplays with the host during infection and how SARS-CoV-2 infection can cause so many disease symptoms. Our analysis of three different datasets (COVID-19, recovered, and healthy) revealed significantly higher DEGs in COVID-19 patients than recovered humans and healthy controls. Some of these DEGs were found to be co-expressed in both COVID-19 patients. They recovered humans supporting the notion that DEGs level is directly correlated with the viral load, disease progression, and different comorbidities. The protein-protein interaction consisting of 24 nodes and 72 edges recognized eight hub-nodes as potential hub-proteins (i.e., RPL4, RPS4X, RPL19, RPS12, RPL19, EIF3E, MT-CYB, and MT-ATP6). Protein–chemical interaction analysis identified three chemical agents (e.g., Valproic Acid, Alfatoxin B1, and Cyclosporine) enriched in COVID-19 patients and recovered individuals. Mental retardation, mental deficiency, intellectual disability, muscle hypotonia, micrognathism, and cleft palate were the significant diseases associated with COVID-19 by sharing DEGs.

## INTRODUCTION

In late December 2019, a novel respiratory disease, now popularly termed as “COVID-19”, caused by severe acute respiratory syndrome coronavirus 2 (SARS-CoV-2), emerged in Wuhan, China [1-3]. Immediately after its first outbreak in China, this fearsome virus has emerged as one of the deadliest human pathogens [4]. Due to its worldwide spread and severity, COVID-19 has been declared a public health emergency of international concern by the World Health Organization (WHO) [5, 6]. As of January 10, 2022, COVID-19 disease affected 217 countries and territories, and more than 308 million cases have been confirmed around the globe, with about 5.6 million deaths [7]. In the early stage of the outbreak, the spectrum of clinical manifestations of COVID-19 ranges from the common cold to respiratory failure depending on the demography and environments [2, 5, 8]. However, recent data show that the clinical episodes of COVID-19 may range from asymptomatic infection to critical illness, with a dysregulated inflammatory response to infection a hallmark of severe cases and life-threatening multi-system organ failure [8-11]. In most cases (∼ 80%), patients exhibit mild symptoms, while the remaining ∼20% may develop severe lung injury and death from respiratory failure [12, 13]. Some of the clinically infected patients may suffer from acute respiratory distress syndrome (ARDS) and multiple organ failure requiring intensive care unit (ICU) facilities for life support and medication [13]. Risk factors for severe SARS-CoV-2 include age, smoking status, ethnicity, and male sex [10, 14]. Notably, the persistence and prognosis of COVID-19 are greatly influenced by the patients’ underlying health conditions and age [9, 15]. Given no effective antiviral treatment and slow vaccine rollout, COVID-19 continues to be a serious threat to public health worldwide [16].

Despite increasing global threats of COVID-19, the host immune response against SARS-CoV-2 infection remains poorly understood, and the perturbations result in a severe outcome [12, 17]. The nasal epithelium serves as a portal for initial infection and transmission of the SARS-CoV-2 [5]. SARS-CoV-2 employs ACE2 (Angiotensin-converting enzyme 2) as a receptor for cellular entry [18], and the binding affinity of the S protein and ACE2 was found to be a major determinant of SARS-CoV-2 replication rate and disease severity [17, 19]. After the entrance into the susceptible host, SARS-CoV-2 infects cells of the respiratory epithelium and mucous membranes, such as those of the nose or eyes [18, 20]. The host immune response to SARS-CoV-2 infection involves activating cellular and humoral arms. The innate immune system recognizes the SARS-CoV-2 RNAs through three major classes of cytoplasmic pattern recognition receptors: Toll-like receptors (TLRs), RIG-I-like receptors (RLRs), and NOD-like receptors (NLRs) [17, 21]. This response involves the release of interferons (IFNs) and inflammatory cytokines, including the IL-1 family, IL-6, and TNF, that activates a local and systemic response to infection [5, 17]. This inflammatory response cascade involves the recruitment, activation, and differentiation of innate and adaptive immune cells, including neutrophils, inflammatory myeloid cells, CD8+T cells, and natural killer (NK) cells [12]. The resolution of infection is largely dependent on the cytotoxic activity of CD8+T cells and NK cells, which enable the clearance of virus-infected cells [5, 17]. It is believed that dysregulated host immune response leads to the persistence of virus-infected cells and may facilitate a hyper-inflammatory state termed Macrophage (MΦ) activation syndrome (MAS) or “cytokine storm”, and ultimately damage to the infected lung [12, 17]. However, the underlying molecular mechanisms of the aberrant inflammatory responses in serology and histopathology under SARS-CoV-2 infection are still not clear.

The ongoing pandemic of SARS-CoV2 and lack of comprehensive knowledge regarding the progression of COVID-19 has constrained our ability to develop effective treatments for infected patients. To obtain a complete understanding of the host response to SARS-CoV2, one means to examine gene expression in relevant tissues. Until now, the scant amount of gene expression profiles are available from patients with COVID-19 and have yielded some insights into the pathogenic processes triggered by infection with SARS-CoV-2 [12, 17, 22]. Transcriptomic analyses of cells upon viral infections are extremely useful to identify the host immune response dynamics and gene regulatory networks [12, 23]. However, because of the limited number of samples and preliminary analysis, a full picture of the biological state of SARS-CoV2-affected tissues has not emerged. To address this, we have employed RNA-Seq techniques to investigate the upper airway (nasopharyngeal tract) gene expression profile in 22 specimens of COVID-19 patients (n = 8), recovered (n = 7), and healthy (n = 7) individuals using several orthogonal bioinformatic tools to provide a complete view of the nature of the COVID-19 inflammatory response and the potential points of therapeutic intervention.

Through differentially expressed genes (DEGs) analyses in these datasets, we identified several genes coding for translational activities (e. g. RPL4, RPS4X, RPL19, RPS12, RPL19, EIF3E), ATP-synthesis (MT-CYB, MT-ATP6), transcription factors (e. g. E2F1, MAX, EGR1, YY1, SRF), hub-proteins (e. g. KIAA0355, DCUN1D3, FEM1C, ARHGEF12, THBS1), and mi-RNA (e. g. hsa-miR-19b, hsa-miR-495, hsa-miR-340, hsa-miR-101, and hsa-miR-19a) evidencing a sustained inflammation and cytokine storm in the COVID-19 patients.

## RESULTS

### Differentially expression and distribution of DEGs

To elucidate whether differentially expressed genes (DEGs) contribute to the SAR-CoV-2 inflammatory response and the potential points of therapeutic intervention, we analyzed 22 RNA-seq data of nasopharyngeal epithelial tissue of COVID-19 patients, recovered humans, and healthy controls. To perform RNA-seq analysis, we retrieved datasets from the National Center for Biotechnology Information (NCBI) that belonged to previously published BioProject under accession number PRJNA720904 (https://www.ncbi.nlm.nih.gov/bioproject). We identified 1960 and 153 gene signatures in COVID-19 patients and recovered human NT epithelial tissues, which were differentially expressed compared with healthy controls. We particularly focused on the dysregulation (up or down-regulation) of the identified DEGs during SARS-CoV-2 pathophysiology and its overlap with the recovered or healthy states of the humans. The volcano plots presented in **Fig. 2** show the DEGs for COVID-19 with the red dots. The number of shared DEGs between COVID-19 and recovered datasets is presented in the Venn diagrams (**Fig. 2** C, D). Thirty-seven shared DEGs were identified between COVID-19 patients and recovered subjects. Of the detected DEGs, 1,510 (77.04%) genes were upregulated (Up) during SARS-CoV-2 pathogenesis, of which 1,489 (98.61%) genes had a sole association with COVID-19 patients. Likewise, 90 (58.82%) genes were upregulated in recovered humans, and of them, 69 (76.67%) genes had a sole association with the recovery phage of SARS-CoV-2 infection (**Fig. 2**C). By comparing the upregulated genes between COVID-19 patients and recovered individuals, we found that 21 genes (i.e., RPL4, MT-ND2, SCD5, MT-CYB, EZR) were shared between the conditions (**Fig. 2**C). On the other hand, 450 (22.96%) and 63 (41.18%) DEGs were downregulated (Down) in COVID-19 patients and recovered subjects, respectively, and of them, only 12 genes (i.e., MAFF, ARHGEF12, DCUN1D3, DR1, MT-CO1.) were found to be shared between COVID-19 and recovered cases (**Fig. 2**D). The DEGs shared between COVID-19 positive and recovered people and their relationships from the perspective of adjusted P-value, and log2 fold-change is presented in the heatmaps, respectively (**Fig. 3** A, B). Finally, 33 common dysregulated (Up or Down) genes were presented in a bubble plot to show relationships with 10 log fold-change values (**Fig. 3**C).

**Fig. 1.**
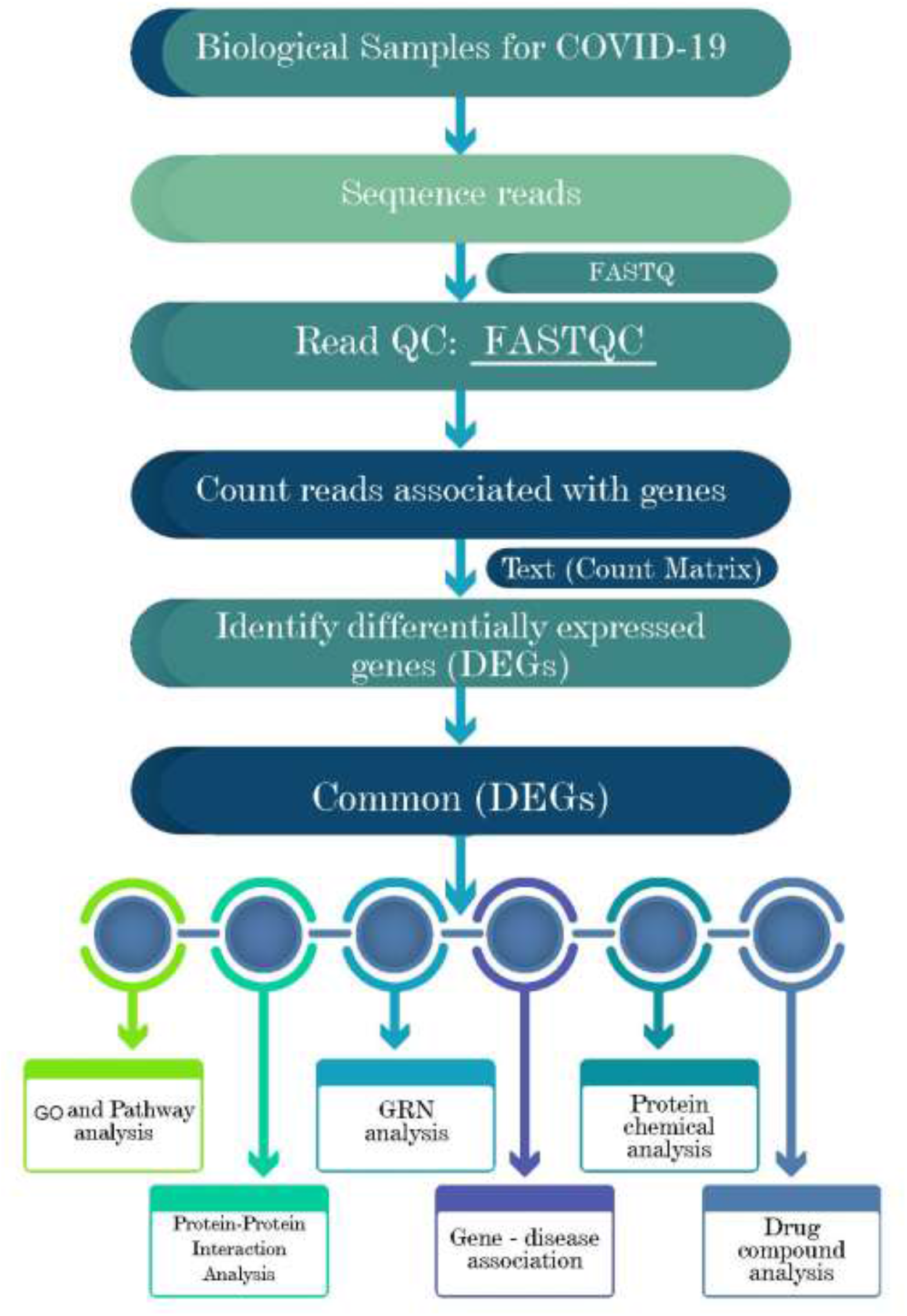
Schematic representations of the paths for differentially expressed genes (DEGs) analysis in RNA-seq data of the COVID-19 patients, recovered humans, and healthy controls nasopharyngeal tract.

**Fig. 2.**
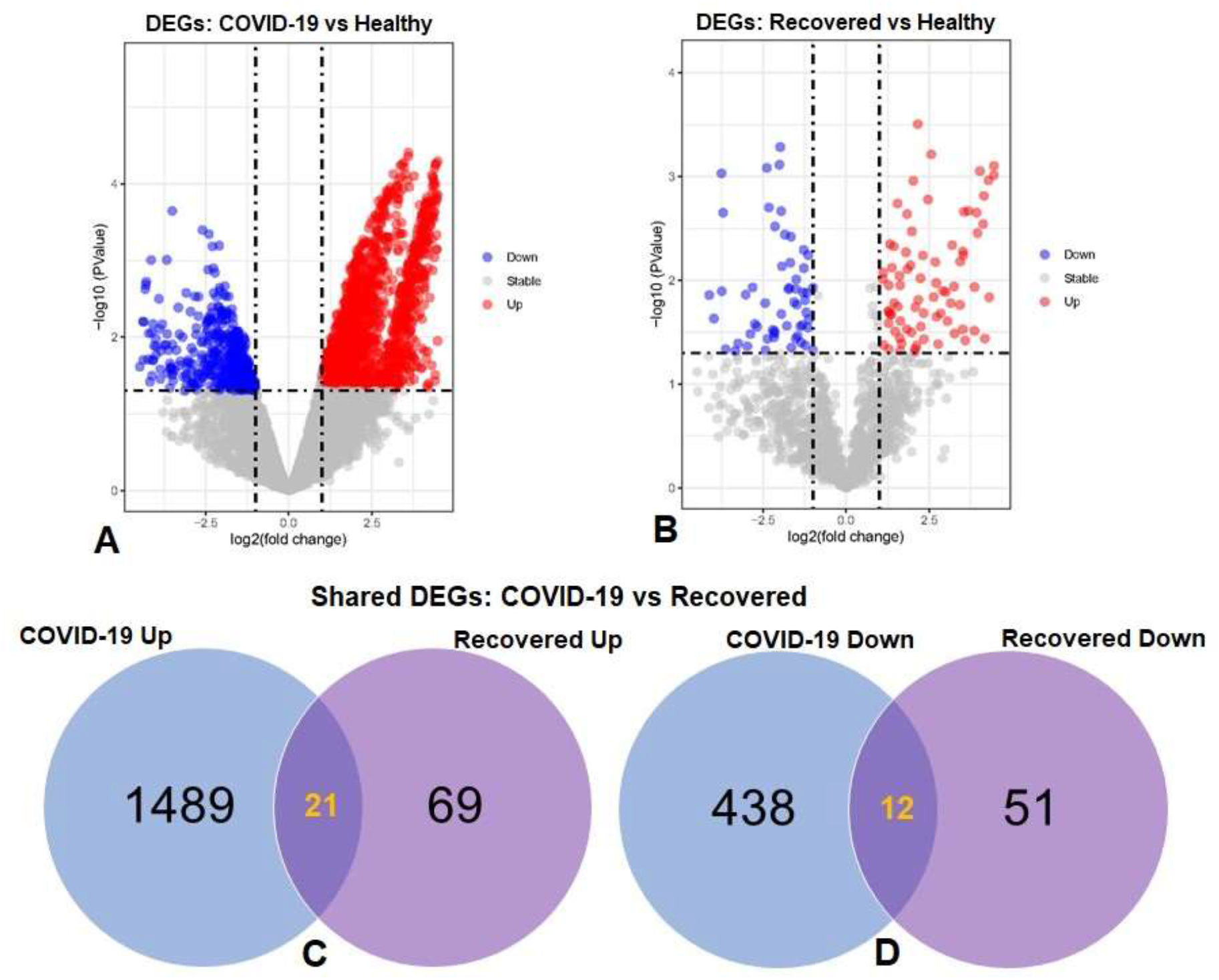
Volcano plots showing dysregulated genes in (A) COVID-19 patients vs. healthy control and (B) recovered humans vs. healthy controls. The red and blue dots indicate the expressions of the upregulated (Up) and down-regulated (Down). Venn diagrams depict the unique and shared DEGs (C) upregulated and (D) down-regulated under the given conditions.

**Fig. 3.**
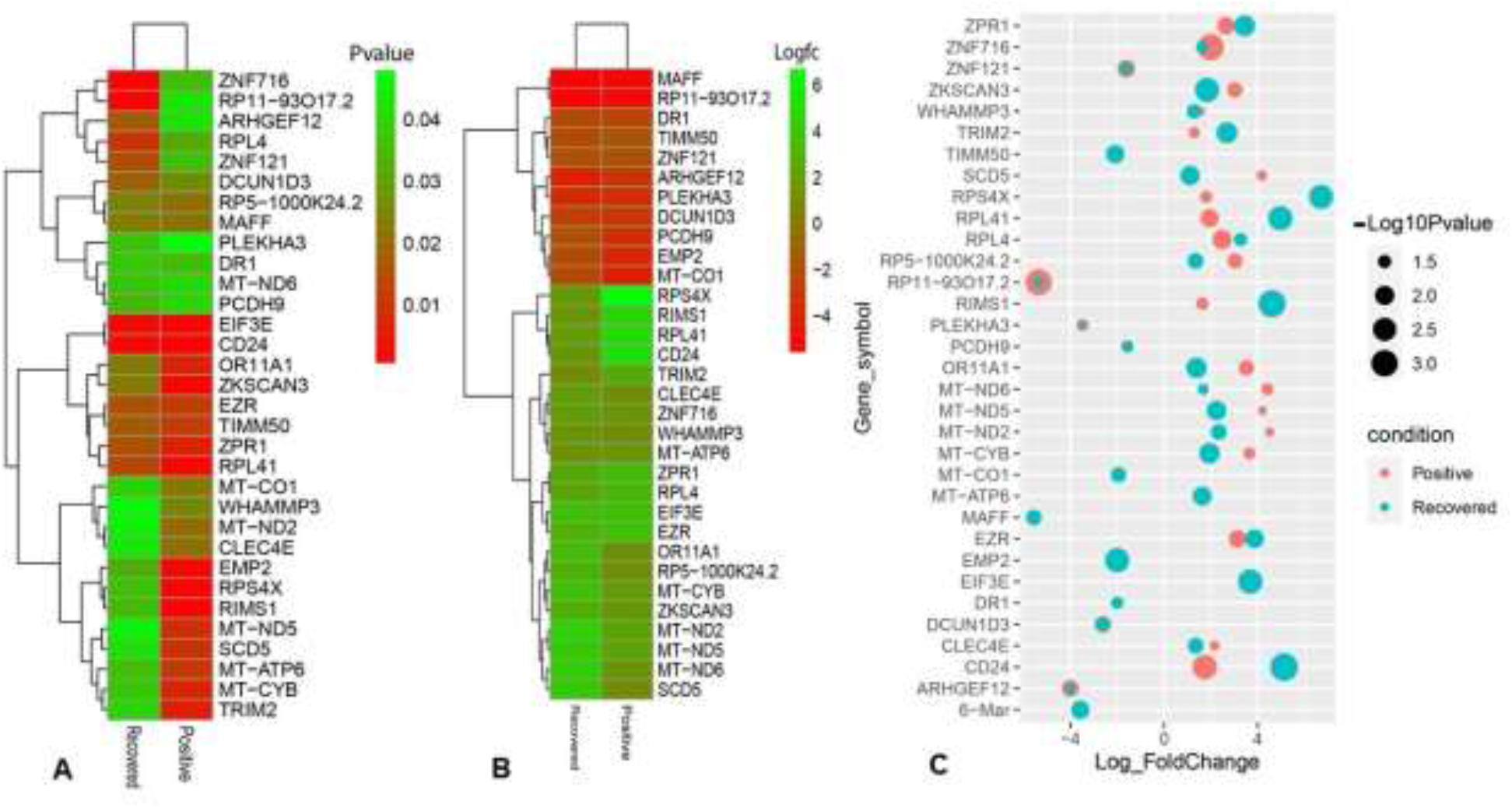
Heatmaps depicting the relationships among common DEGs in COVID-19 patients and recovered subjects based on (A) adjusted P-value and (F) logFC values. (C) Bubble plots showing the combined Log10 fold-changes and p-values for the shared common genes between COVID-19 patients and recovered humans. The red color indicates the genes of COVID-19 patients, and the green color presents the genes of the recovered people, while the mixed color indicates the overlapping genes.

### Functional enrichment analysis identifies significant cell signaling pathways and gene ontology

We used the Enrichr tool to conduct a functional enrichment analysis on the DEGs to identify the signaling pathways and functional gene ontology (GO) keywords significantly enriched with DEGs in the nasopharyngeal epithelial cells from COVID-19 patients. The 33 shared DEGs were used to identify key pathways and GOs that may be linked to COVID-19 comorbidities. We combined all the KEGG and Reactome pathway databases with Enrichr tools to create a single pathway database. We looked at the pathways whose significance was determined by the P-value and plotted the top 20 pathways for each condition (**Fig. 4**). Consideration was given to the paths having a higher logarithmic P-value. The most significant pathways were the ribosome signaling pathway, coronavirus signaling pathway, and c-type-lectin receptor signaling pathway for KEGG analysis (**Fig. 4**B) and forming a pool of free 40S subunits 3-UTR-mediated translational regulation, and eukaryotic translational initiation signaling pathways for the Reactome database (**Fig. 4**B).

**Fig. 4.**
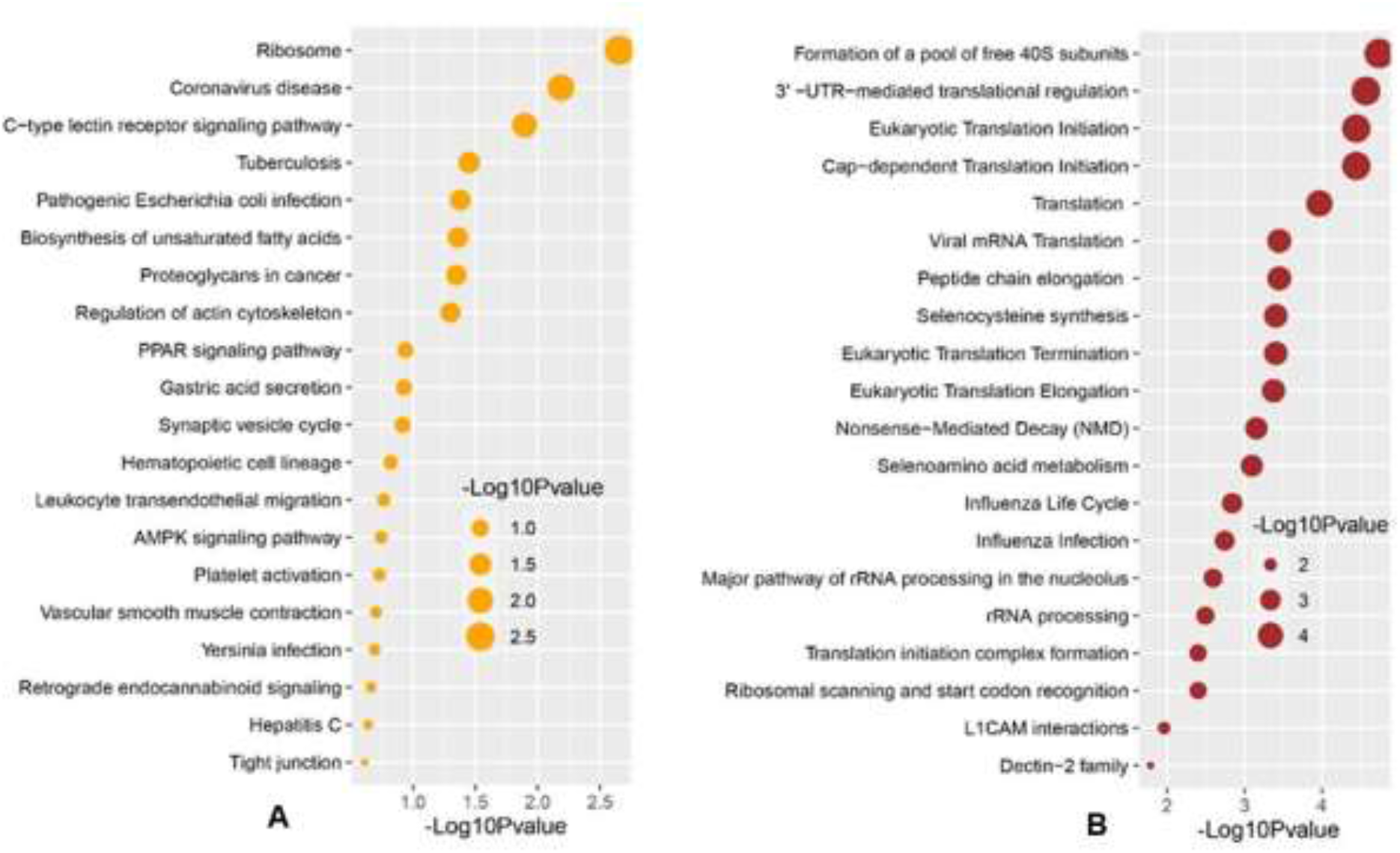
Signaling pathway analysis of nasopharyngeal epithelial cells on COVID-19 patients. We find that the top 20 terms depend on the P-value. (A) KEGG pathway and (B) Reactome pathway.

We used the Enrichr tool to identify significantly enriched cellular signaling pathways and functional GO terms (molecular function, biological process, and cellular component) with DEGs in the nasopharyngeal epithelial cells on COVID-19 patients. The 33 shared DEGs were used to identify key pathways and GOs that may be linked to COVID-19 comorbidities. We looked at the pathways whose significance was determined by the P-value (having a higher logarithmic P-value) and plotted the top 20 pathways for each condition (**Fig. 4**).

Furthermore, we have also conducted GO term enrichment using the same set of common DEGs. We employed the GO biological process, the GO molecular function, and the GO cellular component databases obtained from Enrichr libraries. The significantly enriched GO terms were identified if the enrichment yields the high logarithmic value of the adjusted P-value. The top 20 cellular signaling pathways in the COVID-19 patient’s nasopharyngeal epithelial cells were selected in this study (**Fig. 5**) in relevance to the recovered phase. The most significant GO pathways were the ceramide 1-phosphate transfer activity, and ceramide 1-phosphate binding pathways for the molecular functions (**Fig. 5**A), database nuclear-transcribed mRNA catabolic process and regulation of epithelial cell differentiation pathways for biological process (**Fig. 5**B), and membrane raft, and cytosolic sizeable ribosomal subunit pathways for cellular component (**Fig. 5**C).

**Fig. 5.**
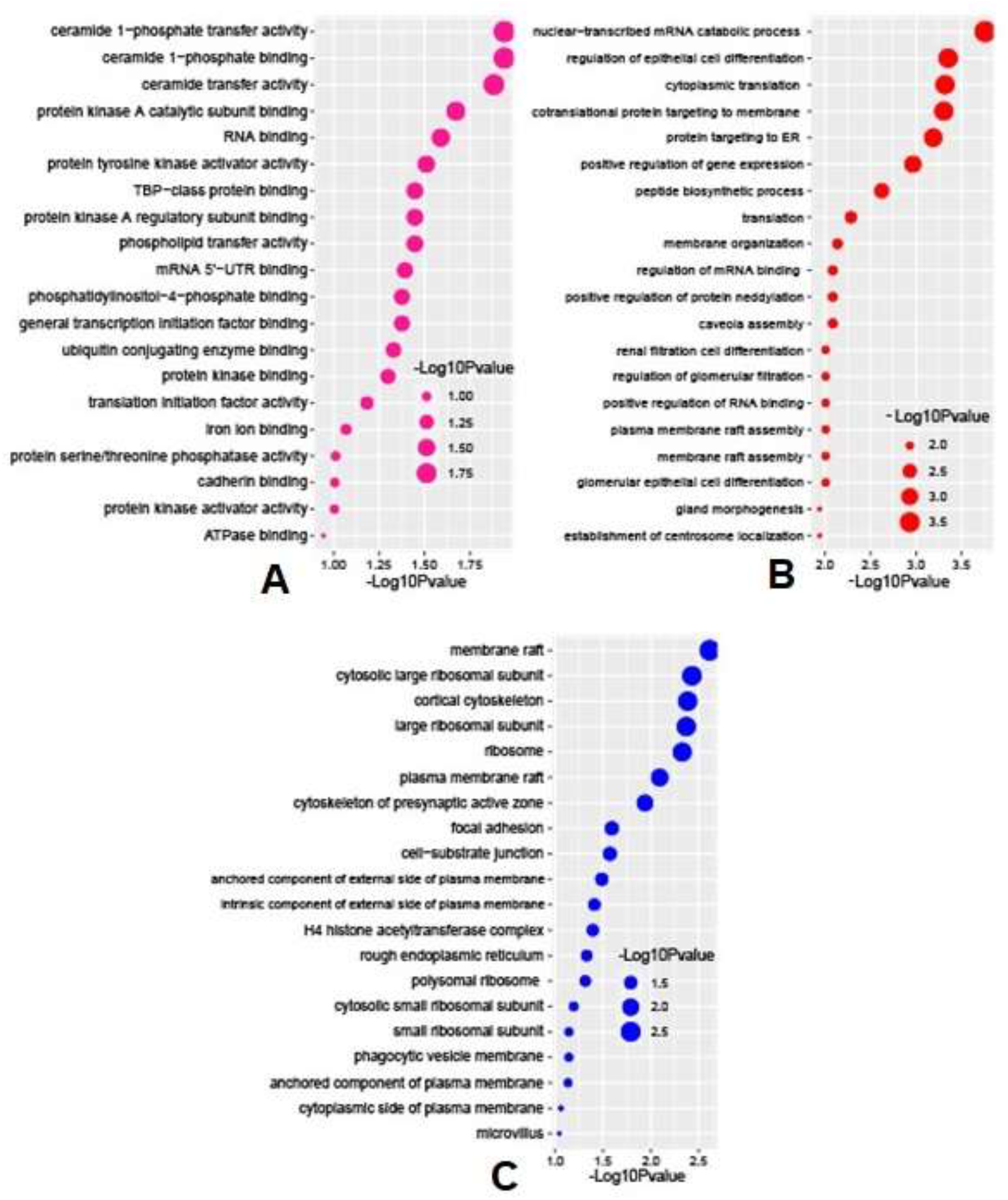
Based on the adjusted P-values, the top 20 cell signaling pathways in the nasopharyngeal epithelial cells on COVID-19 patients. The pathways have been formed by combining the DEGs that are common in the (A) GO molecular function, (B) GO biological process, and (C) GO cellular component.

### Protein-protein interaction network analysis

A Protein-protein interaction (PPI) network was built from the common DEGs interactions, consisting of 24 nodes and 72 edges. The PPI network clustering highlighted RPL4, RPL18A, EIF3E, EIF3D, RPS4X, RPL19, EIF3K, RPS12, MT-ND2, MT-CO1, MT-ATP6, and MT-CYB with high interaction activity (**Fig. 6**). The proteins with several connecting edges can be identified as hub proteins. **Fig. 7** shows the top-10 hub-nodes within the PPI network. As anticipated by five different methods (i.e., maximum neighborhood component; MNC, betweenness, degree, edge percolated component; EPC and maximal clique centrality; MCC), we recognized eight hub-nodes as potential hub-proteins (i.e., RPL4, RPS4X, RPL19, RPS12, RPL19, EIF3E, MT-CYB, and MT-ATP6) (**Fig. 7** A-E). Interestingly, these eight hub proteins were common in all methods. Only RPS4X was found from 4 methods except for betweenness (**Fig. 7**B). Conversely, the betweenness method predicted only three proteins (i.e., SCD5, EZR, and RIMS1) from the shared DEGs as hubs that were not found by other methods (**Fig. 7**B).

**Fig. 6.**
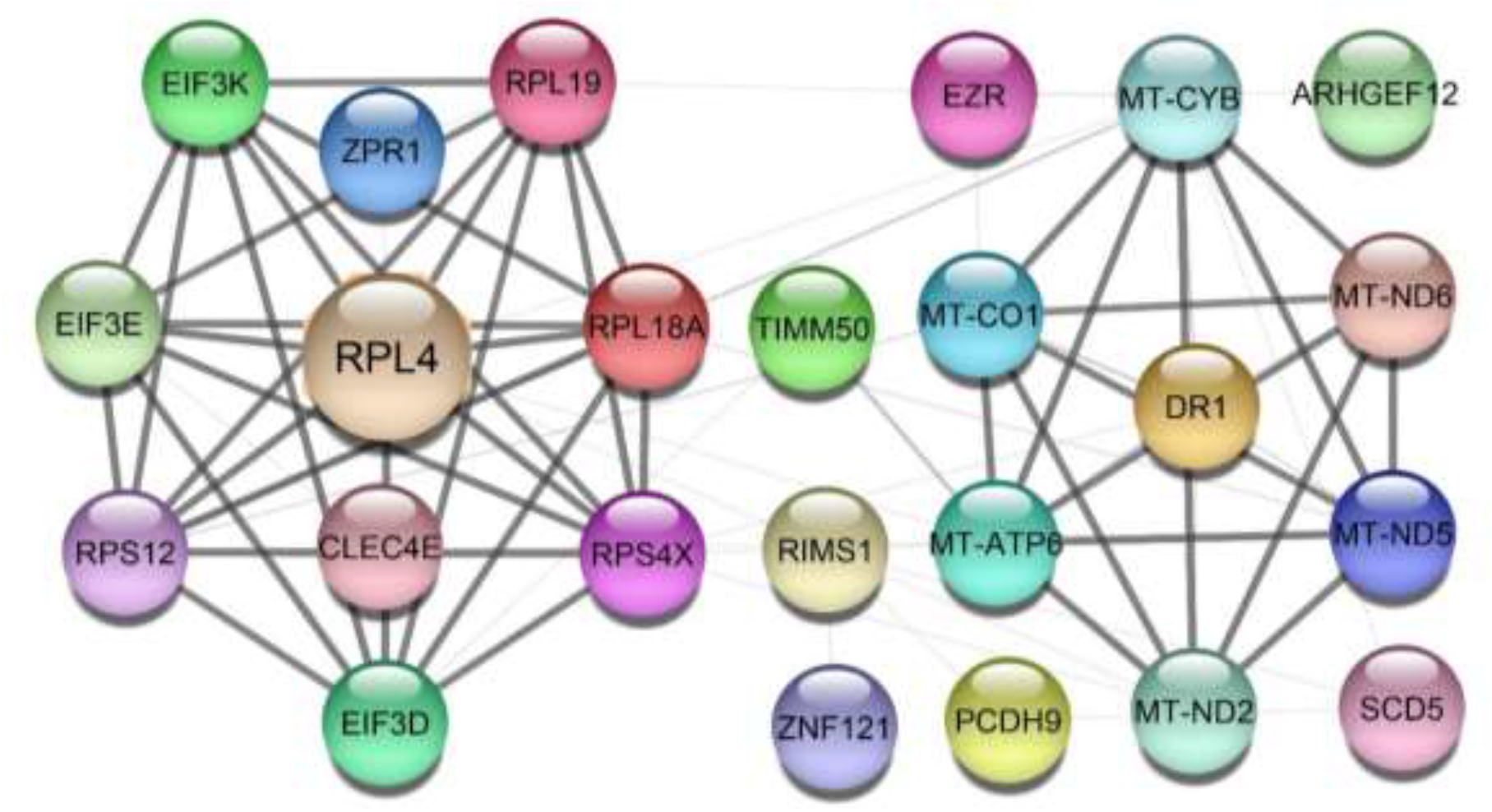
Protein-protein interaction (PPIs) network of common DEGs in COVID-19 patients. The nodes represent the proteins, and the edges represent the interactions across the proteins. Proteins having more edges are highly expressed, and thickness between the edges indicates the strength of interactions.

**Fig. 7.**
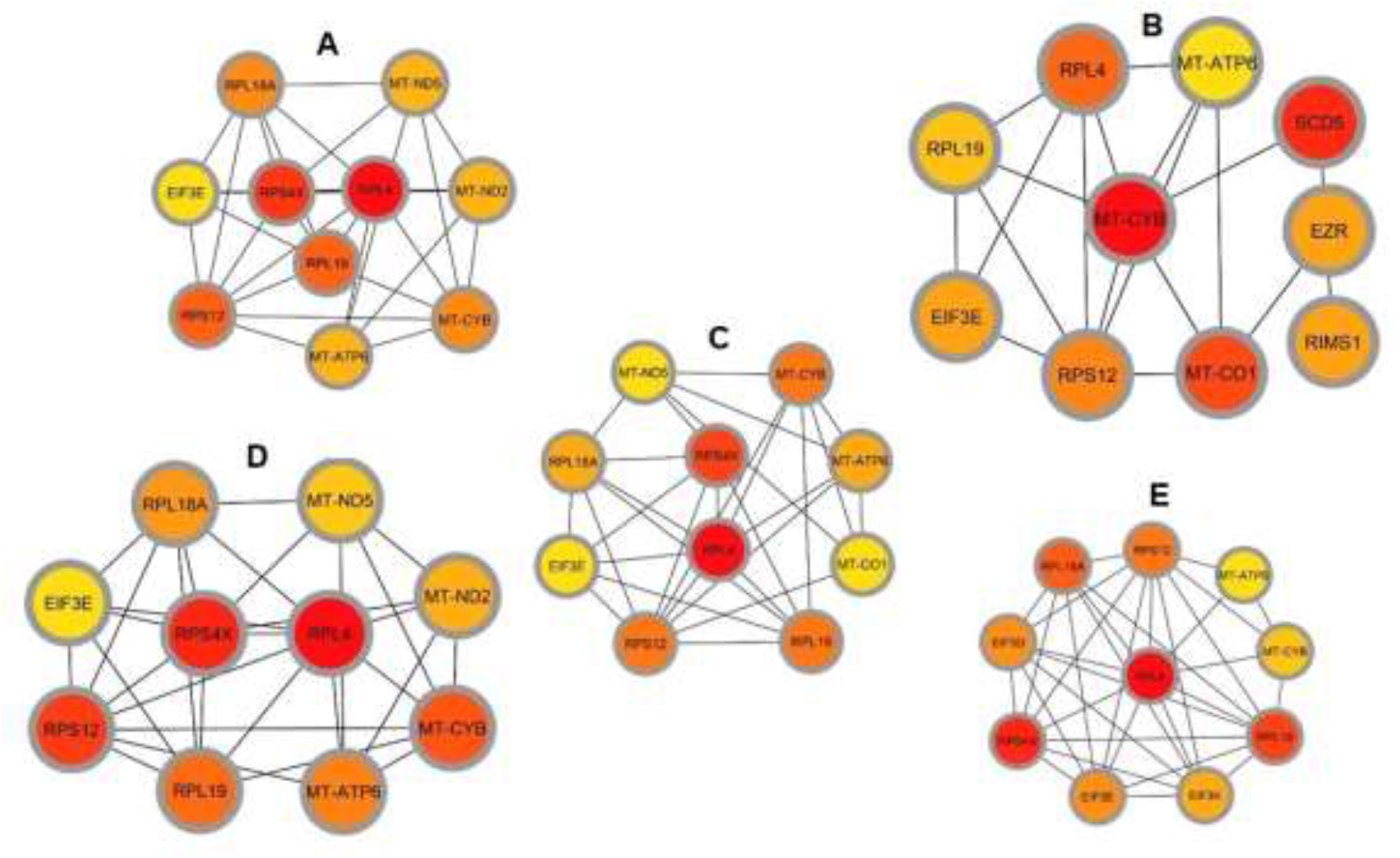
Determination of hub genes from the PPI network by using the Cytohubba plugin in Cytoscape. We applied five algorithms of the Cytohubba plugin to obtain the hub genes. Here (A) maximum neighborhood component (MNC), (B) betweenness, (C) degree, (D) edge percolated component (EPC), and (E) maximal clique centrality (MCC). Red to yellow color gradients indicate the higher ranking of hub genes.

### GRN analysis identifies DEGs–miRNA and transcription factor (TF)–gene interactions

The common DEGs between COVID-19 patients and recovered humans were used in this study. The DEG–miRNA interactions network is depicted in **Fig. 8**A. The dysregulated genes are shown by the circles in the picture, while the squares represent the miRNAs. The association among different nodes of DEGs and miRNA (circles or squares) is represented by different lines linking them. Nodes in a network that connect multiple edges are considered significant nodes since they are more critical. Out of 21 miRNAs detected, hsa-let-7e-5p, hsa-mir-7977, hsa-mir-155-5p, hsa-mir-186-5p, and hsa-mir-1827 were the most expressed miRNAs and had a stronger association with DEGs (Fig. 8A). Likewise, among the DEGs, DMD, AHDC1, BAG4, EMP2, TIMM50, RPL7L1, and THBS1 were more significant since these DEGs have a higher degree (number of connecting edges) among the others and miRNAs (**Fig. 8**A). We further studied the interactions between TF and DEGs and identified 14 TFs, of which FOXC1, FOXL1, NFIC, YY1, and PPARG were significantly enriched and showed more interactions with DEGs (**Fig**. 8B).

**Fig. 8.**
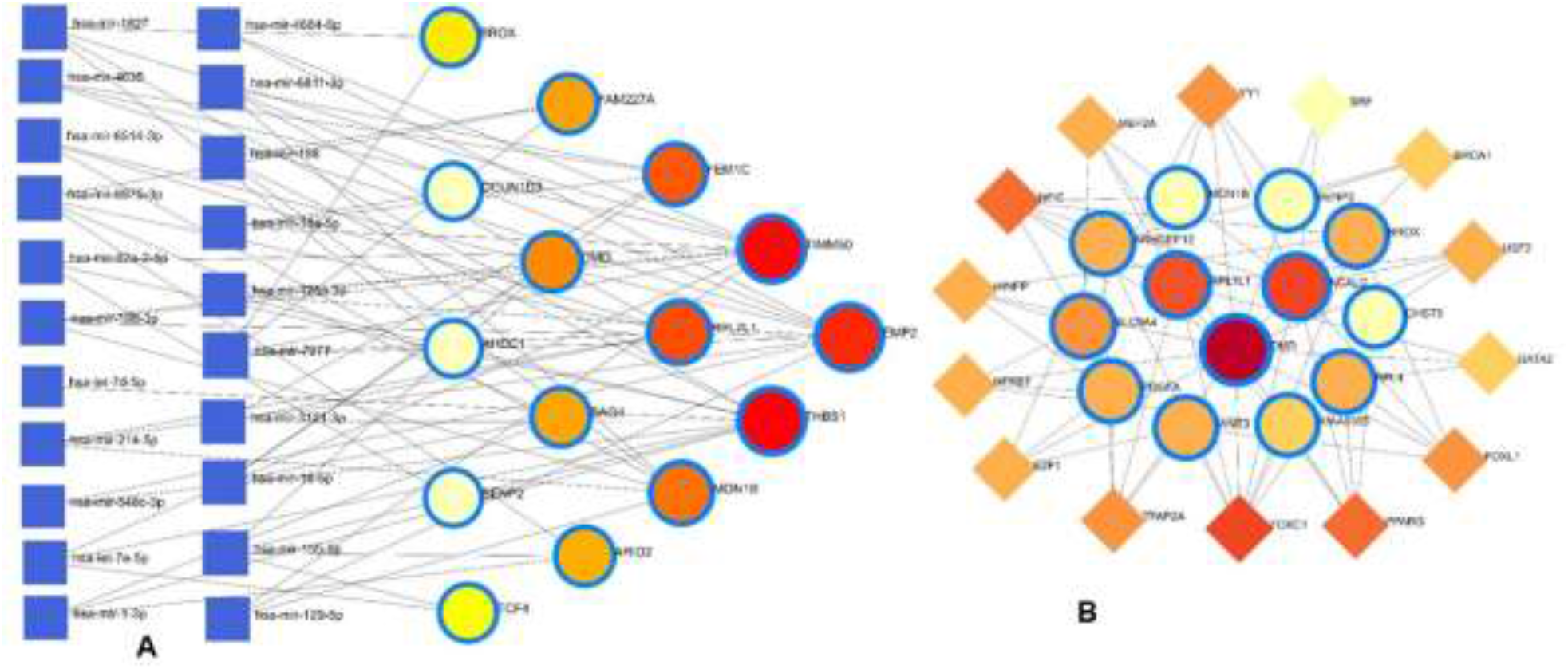
Genes regulatory networks. (A) Gene regulatory networks (gene-miRNA) of the nasopharyngeal epithelial cell on COVID-19 patients with the shared dysregulated genes. The square shapes and circular shapes represent the miRNA and genes, respectively. (B) Gene regulatory networks (transcription factors; TF) of the nasopharyngeal epithelial cell on COVID-19 patients with the shared dysregulated genes. The square shapes and circular shapes represent the TF and genes, respectively.

Apart from these, the present study included TFs and miRNAs that were highly relevant to SARS-CoV-2 interactions. This analysis identified 19 hub proteins, 10 TFs, and 5 miRNAs (**Fig. 9**A). In COVID-19 interaction, the TF–miRNA network showed that E2F1, MAX, EGR1, YY1, and SRF were the highly expressed TFs, and hsa-miR-19b, hsa-miR-495, hsa-miR-340, hsa-miR-101, and hsa-miR-19a were among significant miRNAs (**Fig. 9**A).

**Fig. 9.**
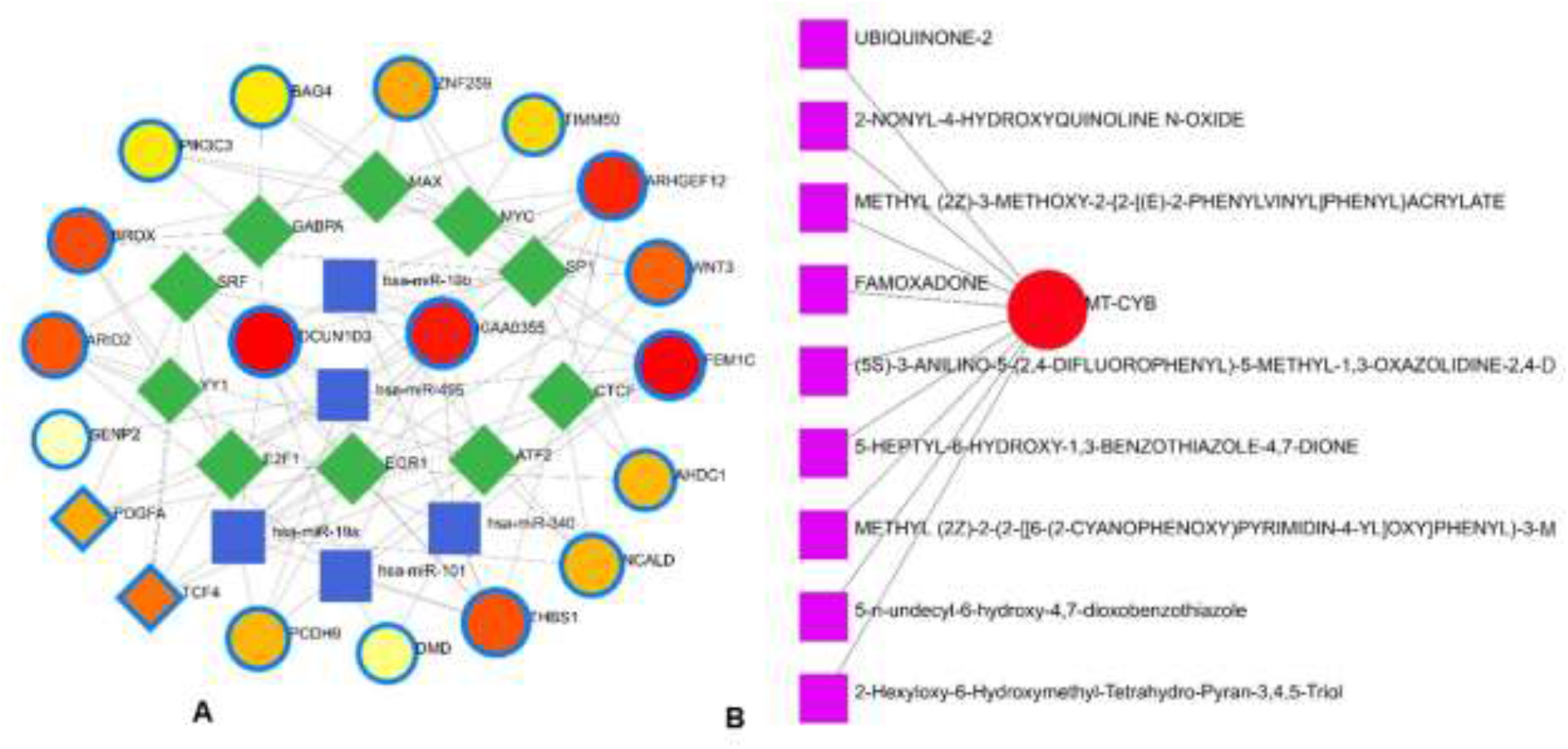
Gene regulatory and protein-drug interactions. (A) Gene regulatory networks (TF-miRNA) of the nasopharyngeal epithelial cell on COVID-19 patients with the shared dysregulated genes. The blue color and square shapes indicate the miRNA, while the green color is square-shapes, representing TF. (B) Protein-drug interaction network. Nine pharmacological compounds are indicated by square shapes (pink color), while one circle shape (red) represents the hub node.

### Protein-drug and protein-chemical interactions reveal possible drugs for COVID-19 patients

Protein-drug interaction (PDI) networks provide a wealth of information about possible pathogenesis mechanisms and drug interactions that may not be evident using conventional approaches. To disrupt the SARS-CoV-2 pervasiveness, we sought to find pharmaceutical compounds that interact with viral proteins (Methods). We detected nine pharmacological compounds (for example, famoxadone, ubiquinone-2, 2-nonyl-4-hydroxyquinoline, 5-n-undecyl-6-hydroxy-4,7-dioxobenzothiazole) acting against one protein, the human mitochondrial cytochrome b (MT-CYB) (**Fig. 9**B).

Protein–chemical interaction (PCI) is an important study to understand the functionality of proteins underpinning the molecular mechanisms within the cell, which may also help in drug discovery. For example, it has been discovered that SARS-CoV-2 infection causes PCI networks in the COVID-19 patients and recovered humans. **Fig. 10**A depicts a network of PCI among significant proteins. The significant proteins identified from this network include FEM1C, NCALD, THBS1, PCDH9, DMD, and PDGFA. Similarly, we identified three chemical agents such as Valproic Acid, Alfatoxin B1, and Cyclosporine enriched in this interaction analysis (**Fig. 10**A).

**Fig. 10.**
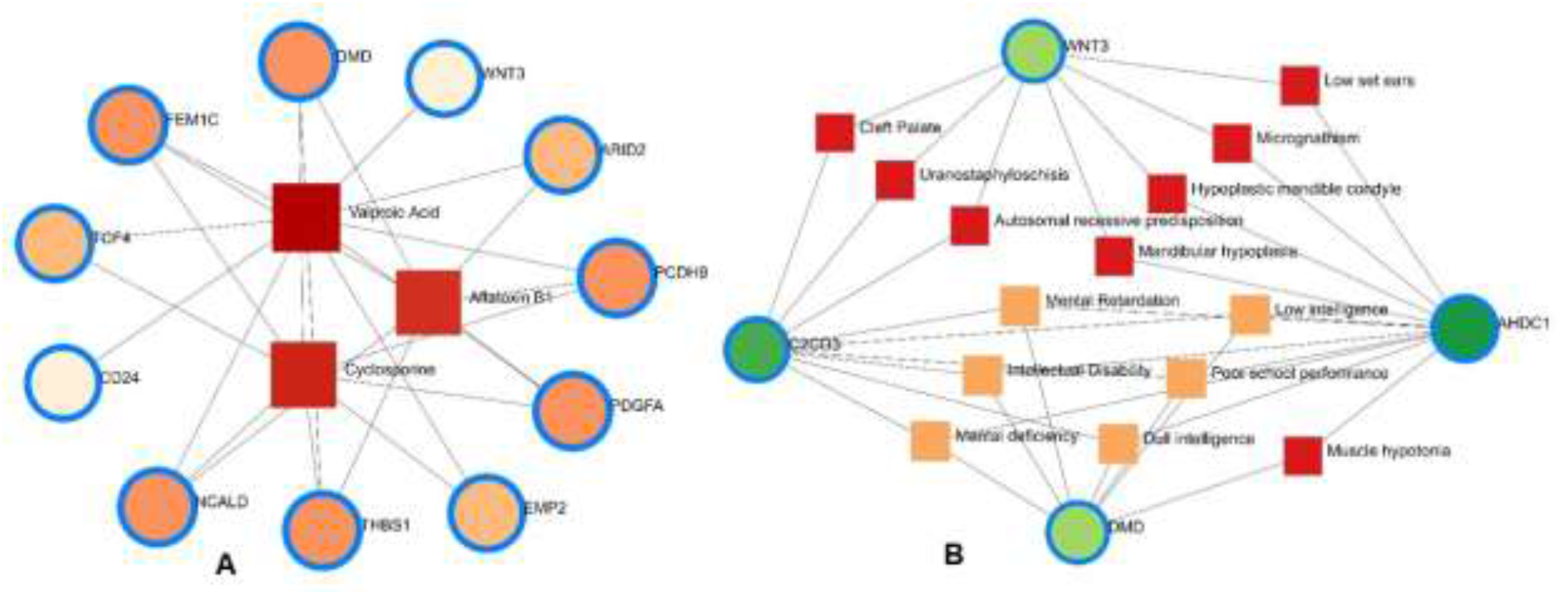
Protein-chemical and gene-disease association. (A) Protein-chemical interaction network. Three phytochemical compounds were found against 11 genes. Circles showed the shared DEGs, while square shapes indicated interacting phytochemical compounds. (B) Gene-disease association network. Circles indicate the common differential genes, and the 14 square shapes represent the common diseases interconnected to the COVID-19 patients.

### Gene-disease network finds different diseases associated with COVID-19

This study hypothesizes that many conditions can be associated or connected with COVID-19 by sharing some common genes among themselves. Disorder-specific therapeutic interface strategies attempt to discover the link between genes and diseases. In this study, we found 14 other diseases associated with COVID-19 by sharing four DEGs (i.e., DMD, C2CD3, WNT3, and AHDC1) most prevalent in COVID-19. Of the detected diseases, mental retardation, mental deficiency, intellectual disability, muscle hypotonia, micrognathism, and cleft palate were the significant diseases interconnected with COVID-19 (**Fig. 10**B).

## DISCUSSION

The SARS-CoV-2 infection causes many illnesses, from minor respiratory sickness that is generally asymptomatic to severe pneumonia that results in multisystem failure and death. The effects of SARS-CoV-2 infection on differential gene expressions (DEGs) in the nasopharyngeal epithelial cells of COVID-19 patients and recovered persons are investigated in this study. We found 1960 and 153 DEGs differently expressed in COVID-19 patients and recovered individuals, respectively, compared to healthy controls. More than 98.0 percent of the upregulated gene signature had a single connection with COVID-19 patients, and 77.0 percent of these DEGs were increased during SARS-CoV-2 pathogenesis. As a result, COVID-19 patients had a higher level of gene upregulation than recovered and healthy persons. As a result, COVID-19 patients had a higher level of gene upregulation than recovered and healthy persons. Previous research has suggested that differences in gene expression across patient groups could be caused by changes in cellular tissue composition, such as the recruitment of immune cell types to the infection site [11]. Jha et al. [24]discovered 338 DEGs in the RNA-seq dataset of lung epithelial cells infected with SARS-CoV-2, including 92 increased and 246 decreased genes across the datasets. In this study, top abundant DEGs such as genes encoding for ribosomal protein (RPL4), controlling the production of the mitochondrial reactive oxygen species (MT-ND2) [25], modulating cell proliferation and differentiation (SCD5) [26], mitochondrial deficiencies, and associated disorders (MT-CYB) [27], epithelial marker ezrin (EZR) associated with cell surface structure adhesion, migration, and organization of the SARS-CoV-2 [28] were found to be co-expressed in the nasopharyngeal epithelial cells of COVID-19 patients and recovered humans (Fig. 2C). Conversely, SARS-CoV-2 infection suppressed the expression genes associated with transcription factors (MAFF) [29], erythropoiesis (ARHGEF12) [30], membrane neddylation (DCUN1D3) [31], global regulator of transcription (DR1) [32], and cytochrome-c oxidase activity (MT-CO1) [33] in both COVID-19 patients and recovered humans (Fig. 2D).

We next investigated whether host gene expression during SARS-CoV-2 pathophysiology is associated with functional enrichment, for example, cell signaling pathways and gene ontology. Our results showed that DEGs related to ribosome signaling pathway, coronavirus signaling pathway, c-type-lectin receptor signaling pathway, forming a pool of free 40S subunits, 3-UTR-mediated translational regulation, and eukaryotic translational initiation signaling pathways were significantly enriched in the nasopharyngeal epithelial cells on COVID-19 patients. These findings corroborated the previously published studies to understand host transcriptional response to influenza A virus and SARS-CoV-2 in primary human bronchial epithelial cells [22, 24]. Gene ontology analysis identified several pathways: ceramide 1-phosphate transfer activity, ceramide 1-phosphate binding pathways, nuclear-transcribed mRNA catabolic process, regulation of epithelial cell differentiation pathways for biological process, membrane raft and cytosolic sizeable ribosomal subunit pathways for cellular component significantly enriched in COVID-19 patients. Ceramide 1 phosphate (C1P) can augment immunity and control COVID-19 infection by enhancing autophagy, adaptive immunity (Th1 programming) MHC-I dependent cytotoxic T lymphocytes (CTL) response [34]. The epithelium lining the airways plays a key role in the defense against infections. Several lines of evidence showed that SARS-CoV-2 infection induces epithelial barrier function, as documented by decreased trans-epithelial resistance, increased permeability, and altered distribution of the tight junction protein [35, 36]. However, this functional impairment remained transient, with signs of epithelial regeneration during the recovery phage of SARS-CoV-2 infection. Basal cell mobilization and replication can also be observed to exert a moderate effect on epithelial barrier integrity [35]. With these dysregulated genes, we have conducted the PPI network analyses. PPIs network analysis is the most prominent section of the study as hub gene detection, analysis of modules, and drug identification thoroughly depends on the PPIs network. According to the PPIs network (Fig.s 6 and 7), ribosomal proteins (RPL4, RPS4X, RPL19, RPS12, RPL19), translation initiation factor 3 subunit E (EIF3E), mitochondrial deficiencies and associated disorders (MT-CYB) [27], and cytochrome-c oxidase activity (MT-CO1) [33], and mitochondrial oxidative phosphorylation (MT-ATP6) proteins were declared as hub proteins because of their high interaction rate or degree value. SARS-CoV-2 infection regulates the mitochondrial transcription of the proteins (MT-CO1 and MT-ATP6) involved in ATP synthesis and respiratory activity, oxidative stress, pro-inflammatory state, and cytokine production [37]. The increased expression of ribosomal proteins can be attributed to the fact that the virus hijacks the host’s translational machinery for its survival by the mechanisms such as ribosome shunting and phosphorylation of ribosomal proteins [24]. These hyper-interactive proteins’ PPI analysis and gene enrichment analysis showed significant biological functions connected to COVID-19 related to cell signaling pathway and the host response to SARS-CoV-2 infections [38]. As discussed earlier, these proteins are involved in several other disorders [22, 37, 38].

We then investigated the protein-protein, gene–miRNA, TF-gene, protein-drug, and protein-chemical interactions of the common DEGs in COVID-19 patients and recovered humans. Our results showed that hsa-miR-19b, hsa-miR-495, hsa-miR-340, hsa-miR-101, and hsa-miR-19a were the mostly expressed miRNAs (Fig. 8A), and E2F1, MAX, EGR1, YY1, and SRF were the highly expressed transcription factors (TFs) (Fig. 8B). While host responses to infection are critical in differential outcomes of SARS-CoV-2 infection, the role of miRNAs in COVID-19 pathogenesis is poorly understood. We observed that most of these miRNAs were strongly upregulated in COVID-19 patients, which could be used as the circulating biomarkers for the diagnosis or prognosis of COVID-19 [39]. Circulating miRNAs are extracellular serum/plasma miRNAs that could be involved in cell-cell communication and thus, might contribute to disease progression. Besides their diagnostic value, miRNAs are well known for their therapeutic potential, especially in viral diseases. A recent report has compared the miRNA signature in the peripheral blood of COVID-19 patients versus healthy donors, and several miRNAs have been identified to be deregulated and interfered with the shaping of the immune responses [40]. Therefore, the upregulated levels of miRNAs could be involved in the inflammatory storm seen in COVID-19 patients via inhibiting the immunosuppressive and anti-inflammatory role ensured by the transcription signaling pathway. Recent studies reported that transcription of mRNAs in epithelial cells is induced by TNF-α and triggers a negative feedback loop involving E-selectin to control inflammatory signaling [24, 41]. Although we identified the 14 TF-genes showing more interactions with DEGs, we further assess whether these genes have the potential causal effects on the COVID-19 development. Our network-based approach identified TF hubs that likely regulate many cellular functions (e.g., cytokine storm) overexpressed i COVID-19 patients. Previous research identified 95 TFs in cytokines upregulated in the COVID-19 patients, and of them, 19 TFs are targets of FDA-approved drugs [42]. Targeting TFs associated with the cytokine releasing syndrome provides candidate drugs and targets to treat COVID-19 [42]. However, additional research is needed to determine whether these combinations elicit the same immunomodulatory response in the context of SARS-CoV-2 infection.

Nine pharmacological compounds were found to be effective against SARS-CoV-2, and of them, fungicide (famoxadone), blood pressure controlling coenzyme (ubiquinone-2), secondary metabolite producing quinoline (2-nonyl-4-hydroxyquinoline), and xenobiotic compound (5-n-undecyl-6-hydroxy-4,7-dioxobenzothiazole) showed their activity against the human mitochondrial cytochrome b (MT-CYB). These compounds have significant antimicrobial, antidiabetic, anti-inflammatory, antiviral, and antioxidant activities [43] against SARS-CoV-2 infections. Furthermore, protein–chemical interaction (PCI) showed that FEM1C, NCALD, THBS1, PCDH9, DMD, and PDGFA proteins showed interactions with three chemical agents such as Valproic acid (VPA), Alfatoxin B1, and Cyclosporine. Numerous promising antiviral therapies against SARS-CoV-2 are being investigated with the hope of preventing both interindividual transmission and severe complications of the COVID-19. The VPA can reduce the SARS-CoV-2 receptor ACE-2 expression level and be used as a potential drug candidate for the prevention strategy against COVID-19 [44]. Aflatoxin B1 (AFB1), which alters immune responses to mammals, is one of the most common mycotoxins in feeds and food and a potential aggravating risk factor in COVID-19 patients [45]. The effect of cyclosporine on coronaviruses, including the new SARS-CoV2, has been extensively studied [46]. Several earlier studies showed that cyclosporine could prevent uncontrolled inflammatory response, SARS-CoV-2 replication, and acute lung injury [46, 47]. Therefore, there is an urgent need for effective drugs targeting this life-threatening complication, particularly for patients developing acute respiratory distress syndrome. In addition, we identified 14 other diseases associated with COVID-19 by sharing four DEGs (i.e., DMD, C2CD3, WNT3, and AHDC1) which were most prevalent in COVID-19. People with SARS-CoV-2 infections often have coexisting conditions like mental retardation, mental deficiency, intellectual disability, muscle hypotonia, micrognathism, and cleft palate. There is a shortage of information regarding the impact of COVID-19 in patients with tuberculosis, HIV, chronic hepatitis, and other concurrent infections. COVID-19 patients developed serious symptoms, including difficulty breathing, chest pain, loss of muscle control, severe inflammation, and organ damage. The adverse health and economic impact of the COVID-19 pandemic influenced mental health, causing distress, anxiety, and depression [48]. These complications are not necessarily short-lived and can cause long-term effects of multi-organ injury following SARS-CoV-2 infections. COVID-19 appears to present a greater risk to people with intellectual and developmental disabilities, especially at younger ages, and recent evidence suggests that mental health problems significantly increased worldwide during this pandemic [49, 50]. Since muscle possesses the ACE2 receptor to which SARS-Cov-2 binds, it follows that the involvement of the muscle could be due not only to the secondary effects of the infection (e.g., reduced oxygen supply from persistent lung disease, perfusion defects from cardiovascular defects, and vascular damage) but also to the direct action of virus (SARS-Cov-2 myositis) [51].

## CONCLUSIONS

Gene expression analysis may potentially reveal disease-pathogenesis pathways and point to novel targets for potential therapeutic approaches. This study examines the RNA-seq data of COVID-19 patients, recovered persons, and healthy individuals to find DEGs and biomarkers between the SARS-CoV-2 pathogenesis and recovery stage from a molecular and cellular standpoint. We found that COVID-19 patients had a much larger number of DEGs than recovered humans and healthy controls and that some of these DEGs were co-expressed in both COVID-19 patients and recovered humans. We used gene expression analysis with the biomarker to identify cellular signaling pathways and GO terms. In the COVID-19 patients, we found several genes coding for translational activities, transcription factors, hub-proteins, and miRNA expressions, all of which indicated a persistent inflammation and cytokine storm. The signaling pathways, GO terms, and chemical compounds discovered in this study could help researchers Fig. out how genes are linked together to find possible therapeutic approaches. However, the DEGs’ direct molecular biological functions and significant pathways discovered in this study should be investigated further to understand better the mechanisms underlying the host response to SARS-CoV-2 and identify potential therapeutic targets and drug candidates for COVID-19.

## MATERIALS AND METHODS

### Overview of the proposed bioinformatics pipelines

Network-based approaches are common to identify and analyze the pathogenesis of SARS-CoV-2. Datasets required in this work were constructed and collected at the initial phase and detailed in the following subsections. Gene expression analysis was performed to identify the DEGs from each dataset (Fig. 1). Next, the common DEGs between two groups of COVID-19 datasets were identified. These common DEGs were further used to discover their protein-protein interactions (PPIs) and to perform gene set enrichment analysis (GSEA) to identify enriched cell signaling pathways and functional gene ontology (GO) terms. Next, the same set of common DEGs was used to discover three types of GRNs: DEGs–micro RNAs (miRNA) network, DEGs– transcription factors (TFs) network, and TF-miRNA network. Finally, protein–chemical compound and protein-drug interactions were also investigated for the common DEGs (Fig. 1).

### RNA-seq data

This study included 22 RNA-seq data of the nasopharyngeal tract (including COVID-19 = 8, recovered = 7, and healthy = 7) to decipher the DEGs associated with SARS-CoV-2 pathophysiology and therapeutic potentials. These data were deposited in the public repository (NCBI) under bioproject accession number <u>PRJNA720904</u> (https://www.ncbi.nlm.nih.gov/bioproject/?term=PRJNA720904) by Hoque et al., 2021 [52]. The demographic information on the study subjects, COVID-19 diagnosis, sample collection, and details about library preparation and RNA sequencing techniques is available in the previously published articles of Hoque et al. [52].

### Dataset preparation and analysis of differentially expressed genes

This study prepared two datasets as COVID-19 patients versus recovered humans with the same healthy control group for analytical purposes. We performed several statistical operations on the datasets to determine the DEGs. Moreover, the Benjamini–Hochberg false discovery rate method was used to provide a good balance between the discovery of statistically significant genes and the limitation of false positives. The BioJupies generator online server (https://maayanlab.cloud/biojupies/) was used for RNA-seq raw data analysis [1]. In this study, genes with adjusted *P*-value < 0.05 and absolute value of log2 fold-change ≥ 1 were considered DEGs. Next, to determine the shared DEGs, we compared two COVID-19 datasets between each other using the Venny v2.1 web tool [2]. In this article, we use the term combined DEGs’ to refer to the collection of these two sets of DEGs, which have been used in the downstream bioinformatics analyses.

### Functional enrichment analysis

We utilized Enrichr [3] with Fisher’s exact test to conduct the functional enrichment analysis with the combined DEGs. After performing an overrepresentation analysis, a collection of enriched cellular signaling pathways and functional GO keywords were discovered, revealing the biological importance of the previously detected DEGs. In Enrichr analysis, we combined the signaling pathways from two libraries, including KEGG and Reactome, to create a single route. Only the important paths for which the *P*-value was less than 0.05 were evaluated and considered after deleting duplicate pathways. For functional GO annotations, we looked at the GO biological process, GO molecular function, and GO cellular component datasets in Enrichr. We selected the most important GO terms based on a set of criteria and with an adjusted *P*-value < 0.05.

### Protein-protein interaction network analysis

The shared DEGs’ protein-protein interaction (PPI) was analyzed using the STRING database [4]. We applied different local- and global-based methods using the cytoHubba plugin [5] in Cytoscape v3.8.2 [6] to determine potential hubs proteins within the PPI network. While the local method ranked hubs based on the relationship between the node and its direct neighbor, the global method ranked hubs based on the interaction between the node and the whole network. In total, five different methods were considered, including three local rank methods, i.e., degree, maximum neighborhood component (MNC), maximal clique centrality (MCC), and two global rank methods, i.e., edge percolated component (EPC) and betweenness. Next, we compared the results and identified the common nodes as the most potential hubs proteins. Finally, the protein networks were analyzed through Cytoscape v3.8.2.

### Differential gene regulatory network (GRN) analysis

The finding of DEG–miRNA, TF–DEG, and TF-miRNA interaction networks is a part of the GRN analysis. Using the Network Analyst platform [7], the commonly dysregulated genes were utilized to identify GRN networks. Discovering DEG–miRNA interaction networks was accomplished by using the miRTarBase [8] database. The JASPAR [9] database was used to identify the TF-DEG interaction network. Employing TF-miRNA coregulatory network database, the TF-miRNA interaction was analyzed. The networks were filtered with a betweenness value of 100 and degree centrality of 0 to 10 to remove unnecessary information.

### Protein-chemical compound analysis

Analyses of protein–chemical compounds can identify the chemical molecules responsible for the interaction of proteins in comorbidities [53]. For example, this study found protein–chemical interactions using the enriched gene (common DEGs) that COVID-19 patients developed several digestive problems. Furthermore, using the Comparative Toxicogenomics Database [10], we have identified the protein– chemical interactions through Network Analyst [7].

### Protein-drug interaction network

One of the key goals of this study is to identify potential therapeutic compounds which could be effective in mitigating SARS-CoV-2 pervasiveness. Using the shared DEGs, we constructed the protein-drug interaction (PDI) network through the NetworkAnalyst v3.0 web server [7] in conjunction with the DrugBank v5.0 database (https://go.drugbank.com/docs/drugbank_v5.0.xsd). We downloaded the network data and conFig.d the data with Cytoscape v3.8.2 to aid the analysis [6].

### Gene-disease association network

DisGeNET (https://www.uniprot.org/database/DB-0218) is a standardized gene-disease association database that incorporates correlations from various sources involving various biological features of disorders. It emphasizes the increasing understanding of human genetic illnesses. We examined the gene-disease connection using a network analyzer [11] to find diseases and chronic problems associated with common DEGs.

## Data availability

All data needed to evaluate the conclusions in the paper are present in the paper. The sequence data reported in this article has been deposited in the National Center for Biotechnology Information (NCBI) under BioProject accession number PRJNA720904.

## Ethics statement

This manuscript utilized RNA-seq data from a publicly available repository, i.e., NCBI, and has no ethical issue to be declared.

